# The dopamine inhibitor GBR12909 improves attention and compulsive behaviour in female rats

**DOI:** 10.1101/2023.07.07.548063

**Authors:** Sara Abdulkader, John Gigg

**Affiliations:** University of Manchester

**Keywords:** ADHD, dopamine, GBR12909, 5C-CPT, Oldham’s method

## Abstract

**Background:** Stimulants such as methylphenidate are the first-line treatment for attention deficit hyperactivity disorder (ADHD). A principal mechanism of action for these drugs is to reduce dopamine reuptake in the striatum. However, the ensuing risk of abuse with such stimulants means there is an urgent need for new, low-risk therapeutic agents. GBR12909 is a highly selective dopamine reuptake inhibitor, making it an important experimental tool. Indeed, this drug completed Phase II clinical trials for treatment of cocaine abuse. Understanding such drugs has the potential to expand our understanding of the striatal dopamine contribution to impulsivity, attention and compulsive behaviour and will help the development of novel targeted treatments for ADHD without an abuse risk.

**Aims:** The aim of this work was to examine the role of low doses of GBR12909 on attention, impulsivity and compulsive behaviour as measured by 5C-CPT. Oldham’s methods was used to determine the presence of a rate-dependent effect.

**Methods:** Female Lister hooded rats were trained to criterion in the 5C-CPT (>70% accuracy, < 30% omission and < 40% false alarms). Effects of GBR12909 (0.25-1 mg/kg) were investigated on attention, impulsivity and compulsive behaviour under challenging task conditions.

**Results:** The two lowest doses of GBR12909 improved selective attention in a rate-dependent manner while the highest dose of GBR 12909 showed a tendency toward improving compulsive behaviour in a baseline-dependent manner. However, GBR 12909 impaired waiting impulsivity in a baseline-dependent manner.

**Conclusions:** GBR12909 has a clearly beneficial effect on attention and compulsive behaviour in the female rat. These findings support further clinical investigation of GBR-type dopamine uptake blockers or GBR analogues to facilitate the discovery of medications for inattentive ADHD, stimulant abuse, compulsive drug seeking behaviour and obsessive-compulsive disorder.

## Introduction

Attention deficit hyperactivity disorder (ADHD) is a psychiatric disorder characterized by three main behavioural deficits: lack of attention, hyperactivity and impulsivity (Mueller et al., 2017). These are observed during early childhood and often persist into adult life. Indeed, children diagnosed with ADHD have a higher risk of developing social and cognitive problems in later life (Lara et al., 2009). The prevalence of ADHD is between 2-7%: 5% of children have difficulties with inattention and hyperactivity and 2-4% of adults are diagnosed with ADHD (Sayal et al., 2018). With increasing age, inattention becomes more prominent while impulsivity diminishes (Faraone et al., 2015; Telzer et al., 2016). Depending on the main symptoms, ADHD can be divided into three categories: lack of attention (most prevalent among girls), hyperactivity (more prevalent among boys) and a third subtype that is a combination of the first two (Rasmussen and Gillberg, 2000). ADHD is more common in boys than in girls. However, ADHD is unrecognised and underdiagnosed in many countries. In the UK, less than 50% of children with inattention and hyperactivity have been diagnosed with ADHD (Faraone et al., 2015). Girls may go underdiagnosed because girls tend to have inattentive traits (less noticeable symptoms) rather than impulsive traits (Faraone et al., 2015). In addition, coexisting anxiety and depression in girls with ADHD may lead to missed diagnoses (Quinn & Madhoo, 2014).

Stimulants such as methylphenidate and amphetamine are the most common pharmacological treatments for ADHD (Mueller et al., 2017). However, these have the potential to lead to abuse and dependence (Ahn et al., 2013). For example, stimulants (‘smart pills’) are commonly used by students to enhance academic performance (Sharma et al., 2014). Non-stimulants (such as atomoxetine and guanfacine) are the second-line treatment for ADHD; however, these are less effective than stimulants and have a delayed onset of action. Therefore, there is a pressing need for new therapeutic agents that act without this dependence potential (Sharma et al., 2014). Many theories have been developed to explain the behavioural deficit in ADHD, with the most widely accepted mechanism being a deficit in dopaminergic neurotransmission as standard ADHD medications are indirect acting dopaminergic receptor agonist (Faraone et al., 2015). Researchers have focused on dopamine because in healthy individuals it has a very important role in attention, behaviour and reward (Krause et al., 2000; Volkow et al., 2007; Bush et al., 2010; Sharma et al., 2014). The behavioural deficits in ADHD have been linked with hypofunction of the catecholaminergic system in brain regions associated with inattention and impulsivity (Mueller et al., 2017). Levels of synaptic dopamine are regulated by dopamine transporters (DAT) and it is thought that upregulation of the latter results in deficits in selective attention and motivation in patients with ADHD through decreased synaptic dopamine levels in brain areas such as the prefrontal cortex and striatum (Volkow et al., 2007; Rubia et al., 2011).

The continuous performance test (CPT) is the standard clinical test for ADHD (Rosvold et al., 1956; Albrecht et al., 2015). The rodent 5-choice version of CPT (5C-CPT) measures attention and impulsivity in a manner consistent with human CPT (Young et al., 2009; Barnes et al., 2012; Van Enkhuizen et al., 2014; Tomlinson et al., 2014; 2015; Hayward et al., 2016). Indirect dopaminergic agonists such as amphetamine, methylphenidate and modafinil are widely used clinically to treat cognitive deficits in ADHD (Faraone et al., 2015) and these drugs have been used in preclinical studies to test predictive validity of the 5C-CPT (Grottick and Higgins, 2002; Tomlinson et al., 2014; Turner et al., 2017; MacQueen et al., 2018; Caballero-Puntiverio et al., 2019). Recently, it has been reported that touchscreen 5C-CPT can reliably reveal a cognitive enhancing effect of amphetamine in mice and human subjects, as systemic administration of amphetamine (a potent dopamine and noradrenaline reuptake inhibitor) under baseline (standard task demand) conditions improved vigilance by increasing hit rate (probability of correct response) without affecting response inhibition (probability of false alarm) and waiting impulsivity (premature response) in both species (MacQueen et al., 2018). Subsequent studies showed that 5C-CPT can reveal a cognitive enhancing effect of methylphenidate or amphetamine in rats, highlighting the translational validity of this task in preclinical studies (Tomlinson et al., 2014; Young et al., 2020). Furthermore, Modafinil, a weak dopamine and noradrenaline reuptake inhibitor, improved 5C-CPT performance in human subjects by increasing hit rate (Cope and Young, 2017). The effectiveness of standard ADHD medications in reversing attentional deficits in ADHD patients supports the importance of DAT inhibitors in improving attentional processes.

Recent pre-clinical studies have focused on separating rats into high-performing and low-performing subgroups based on their baseline performance (rather than combining animals together) to investigate the effect of dopaminergic manipulation on attentional performance in each sub-group under challenging task conditions (Tomlinson et al., 2015; Hayward et al., 2016). Rate dependence is a well-supported phenomenon in behavioural pharmacological studies, in which the effectiveness of pharmacological intervention changes in a manner consistent with baseline performance (Perkins. 1999). The most common type of rate-dependent function is an inverse relationship between baseline performance and the behavioural change following drug administration (Perkins et al., 1999; Bickel et al., 2016). This type of relationship can be divided into three types: (1) full-range effect, in which low baseline values increase and high baseline values decrease; (2) limited low-range effect, in which low baseline values increase with small changes in high baseline values; (3) limited high-range effect, in which high baseline values decrease with small change in low baseline values (Bickel et al., 2016). A less common type of rate-dependent relationship is a direct relationship between baseline and the change following intervention (Perkins et al., 1999)-Analysis of rate-dependence is often used as a last resort when experimenters find no change in group performance. One of the reasons for this could be the statistical analysis used to determine the presence of a rate-dependent effect (Chiolero et al., 2013). Traditionally, baseline dependency is determined by plotting baseline performance against the change following drug administration (‘Simple’ method). However, this type of relationship is affected by regression to the mean and mathematical coupling (Chiolero et al., 2013). To control for the latter, the ‘Oldham’ method (Oldham, 1962) has been used to determine the presence of a rate-dependent effect (Bickel et al., 2016; Snider et al., 2016; Snider et al., 2018). Determining rate dependence is important as it helps to reveal a relationship that would have otherwise gone unnoticed. An example of a clinical study that was re-analysed by Bickel and colleagues (2016) was by Zack and Poulus (2009). Here, the low-impulsive group showed an increased stop signal response time (SSRT) following modafinil treatment while the high-impulsive group showed a decreased SSRT (Zack & Poulus, 2009). Re-analysing the data using individual scores showed an inverse relationship between placebo baseline SSRT and the proportion of change in SSRT following modafinil administration, that is, rate-dependence (Bickel et al., 2016).

Understanding the effects of selective DAT inhibitors on attention and impulsivity in a task with high translational validity may help the development of novel targeted treatments without an abuse and dependence risk. Contrary to other indirectly acting dopaminergic agonists such as amphetamine and methylphenidate, GBR12909 is a highly selective dopamine reuptake inhibitor, making it an interesting experimental tool (Espana et al., 2008; Ahn et al., 2013). It would be insightful to investigate the effect of a selective DAT inhibitor on a task with high translational validity to humans, such as 5C-CPT (Barnes et al., 2012; Tomlinson et al., 2014), to determine whether improvement in signal detection is due to improved attention or response inhibition and if there is a correlation between baseline 5C-CPT measures and magnitude of change after drug administration. Understanding the contribution of DAT to waiting impulsivity and selective attention may help the development of selective therapeutic agents. For this reason, we investigated the role of a selective DAT inhibitor in modulating attentional processes and inhibitory response control as measured by 5C-CPT. As the Oldham method helps in revealing baseline-dependent effects of stimulants on waiting impulsivity (Bickel et al., 2016) we used correlational analyses to account for natural variation in baseline data when evaluating the effectiveness of GBR 12909 on 5C-CPT performance measures. We compared the traditional method of presenting data with correlational analysis using Oldham’s method to determine if this drug modulates performance measures in a rate-dependent manner.

## Materials and Methods

### Animals

Female Lister hooded rats (n=22; Charles River, UK) were housed in groups of 5 in a temperature (21 ± 2 °C) and humidity (55±5%) controlled environment (University of Manchester BSF facility). Rats weighed 230±10g upon arrival and were allowed free access to food (Special Diet Services, UK) and water for one week prior to the beginning of training. Home cages were individually ventilated cages with two levels (GR1800 Double-Decker Cage, Techniplast, UK) and testing was completed under a standard 12-hour light: dark cycle (lights on at 7:00 am). Two days before training commenced food restriction was initiated to encourage their task engagement and this continued throughout training. During training rats were maintained at approximately 90% of their free-feeding body weight (fed 10 g rat chow/rat/day) and were provided with free access to water. All procedures were conducted in accordance with the UK animals (Scientific Procedures) 1986 Act and local University ethical guidelines.

### Rat 5C-CPT Apparatus

Training took place in 8 operant chambers (Campden instruments Ltd, UK), each placed inside a ventilated and sound-attenuating box. The chambers were of aluminium construction 25cm × 25 cm. Each chamber had a curved wall containing nine apertures. For 5C-CPT training, four of the apertures were blocked by metal caps while the other five were left open (numbers 1, 3, 5, 7 and 9). Each aperture had a light and an infrared beam crossing its entrance to record beam breaks following nose poke. A reward dispenser was located outside each chamber to automatically dispense reinforcers (45 mg sucrose food pellets; Rodent Pellet, Sandown Scientific, UK) into a food tray at the front of the chamber. Entrance to each food tray was covered by a hinged panel so that tray entries were recorded when the rat’s snout opened the panel. House lights were placed in the side wall of the chambers and performance was monitored by a camera attached to the roof.

### 5C-CPT behavioural training

The 5C-CPT benefits from non-target trials in which all five holes are lit and the rat must not respond at any of the five positions to be rewarded (Young et al., 2009; Barnes et al., 2012). The incorporation of non-target trials helps in dissociating response inhibition from waiting impulsivity; premature responses during target trials reflect a deficit in waiting impulsivity, while the failure to withhold responding (false alarms) during non-target trials represents response disinhibition (Young et al., 2011; Barnes et al., 2012). This task also allows the assessment of vigilance in rodents in a manner consistent with human CPT using signal detection theory, facilitating translation of rodent outcomes to the clinic (Young et al., 2013). Training was conducted in a similar manner to previous studies using 5C-CPT (Tomlinson et al., 2014).

### Signal detection theory (SDT)

Two primary parameters are calculated from 5C-CPT trial-outcome measures. The first is hit rate and the second is false alarm rate (Young et al., 2011; Barnes et al., 2012). From these two parameters, sensitivity (vigilance) and bias can be calculated using SDT (see Table 1). Sensitivity reflects the ability to respond to the illuminated hole and inhibit that prepotent response when all five holes are illuminated. Sensitivity can also be measured non-parametrically by calculating the sensitivity Index (SI) (Young et al., 2011; Barnes et al., 2012). 5C-CPT performance can also be affected by bias (tendency to respond to stimuli), which can be measured by calculating the responsivity index (RI) (Young et al., 2013).

**Table 1.** Parameters generated from 5C-CPT trials. Hit = correct response, Miss = omission, CR = correct rejection, FA = false alarm, FAR = false alarm rate, HR = hit rate, *d*’ = discriminability index, SI= sensitivity index and RI = responsivity index.

| Trial Type | Measure | Calculations | SDT parameters |
| --- | --- | --- | --- |
| Target trials | Hit and Miss | $HR = \frac{Hit}{Hit + Miss}$ | $SI = \frac{HR - FAR}{2(HR + FAR) - (HR + FAR)^2}$ |
| Non-target trials | CR and FA | $FAR = \frac{FA}{FA + CR}$ | $RI = \frac{HR + FAR - 1}{1 - (FAR - HR)^2}$ |

### Experimental Design

#### Drug preparation

GBR 12909 dihydrochloride was purchased from Sigma (UK) and dissolved in distilled water. Drug was freshly prepared each day before testing and was injected via the intraperitoneal (i.p.) route 20 min before behavioural testing at a dose volume of 1 ml/kg. All drug doses were calculated as base equivalent weight. The doses used in this study were chosen based on previous studies in rats utilising 5-CSRTT (Fernando et al., 2012).

#### Behavioural testing

Twenty-two rats were used to investigate the effect of GBR12909 on 5C-CPT performance using challenging task conditions that consisted of a variable inter-trial interval (8, 9, 10, 11 and 12 s) and a variable stimulus duration (SD: 1, 0.75 and 0.5 s). The session lasted no longer than 60 min. The inter-trial interval (ITI) was increased to provoke impulsive behaviour. Using a within-subject design rats received 0.25, 0.5, 1mg/kg GBR or vehicle, 20 min before the task. All animals received a standard training session between the testing sessions to ensure performance was maintained at a stable baseline. Each test day was followed by at least a 7-day washout period, after which behavioural testing re-commenced. Prior to the first test day, all animals had been habituated twice to i.p. saline injections.

#### Statistical analysis

All data are displayed as observed mean ± SEM. Overall performance was analysed by one-way repeated measures ANOVA followed by Planned Comparisons. Analysis of the performance across the duration of the session was carried out by two-way repeated measures ANOVA (Trial and treatment as within-subject factors) followed by Planned Comparison on the predicted means. If the outcome of the repeated-measures ANOVA yielded significant effects of dose, further post-hoc analysis was performed using Dunnett multiple comparison test. Alpha level was set to 0.05 and all analysis was carried out using GraphPad Prism (v9.0).

#### Correlational analysis

This study investigated if baseline performance predicted the degree of change in performance following administration of GBR using Oldham method. Oldham’s method was used to determine the presence of a rate dependent relationship. Pearson’s correlation was used when individuals were normally distributed. Nonparametric Spearman’s correlation was used when individuals were not normally distributed. Oldham’s *r* > 0.3 indicates the presence of a rate dependent relationship (Oldham, 1962; Bickel et al., 2016; Snider et al., 2016; Snider et al., 2018). An outlier may increase the value of *r*, therefore, Grubbs test was used to detect an outlier.

## Results

### Effect of GBR 12909 on waiting impulsivity as measured by 5C-CPT performance

The main effect of GBR 12909 on motoric impulsivity averaged across different ITI is shown in Fig. 1A. GBR 12909 showed a tendency towards increasing premature responding. One-way repeated measures ANOVA revealed no significant treatment effect [F (2.712, 56.95) = 2.320, P=0.0908]. Correlational analysis is shown in Fig. 1B. The Oldham method revealed a positive association between baseline impulsivity and magnitude of change following administration of GBR, indicative of a rate dependent effect.

**Figure 1.**
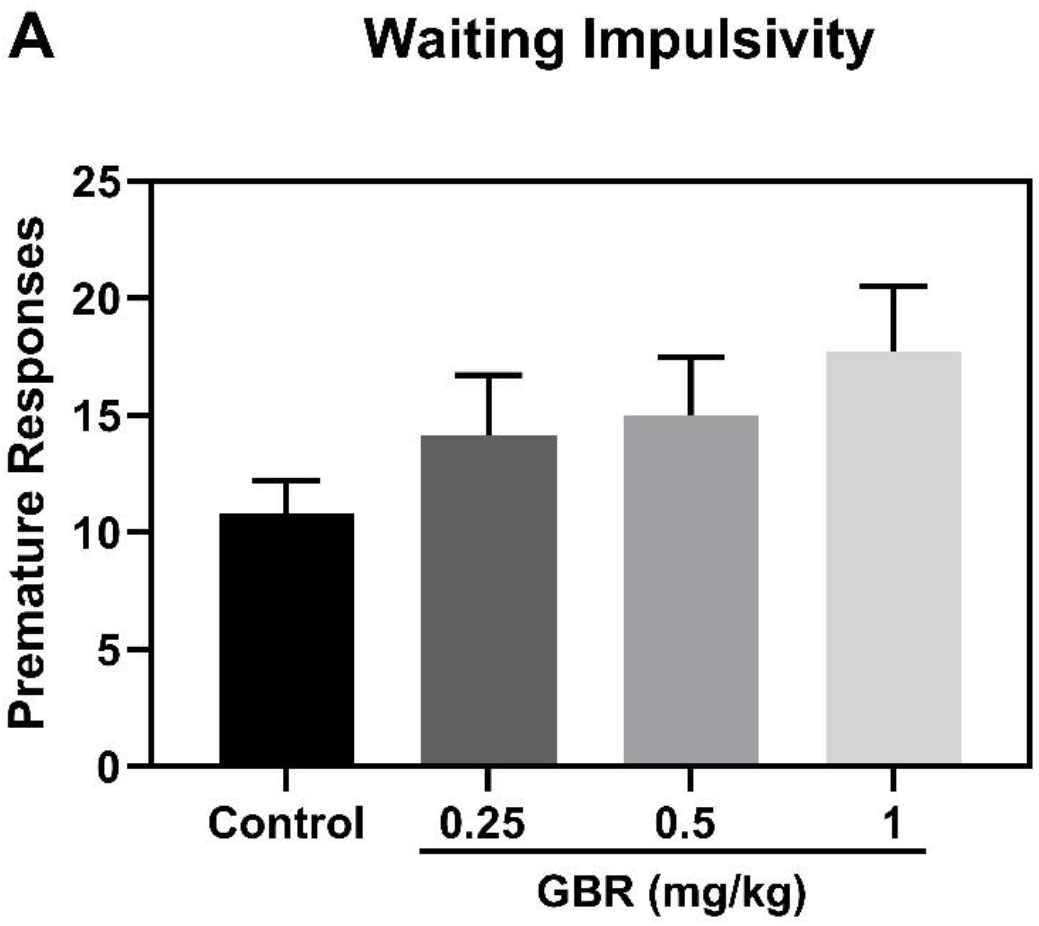
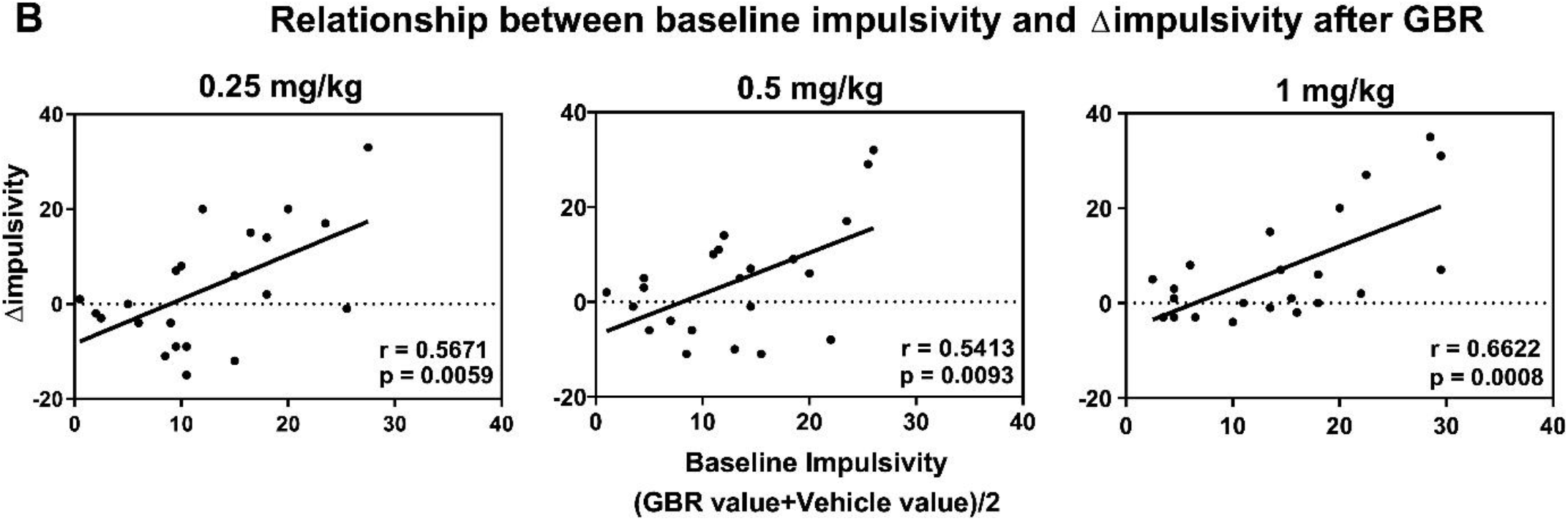
Effects of GBR 12909 (0.25, 0.5 or 1 mg/kg) on waiting impulsivity of rats tested under variable ITIs and variable SDs (n=22). (A) Group impulsivity performance. (B) Correlational analysis (Oldham’s method). Measures are shown as observed mean ±SEM.

### Within-session performance (waiting impulsivity and motivation)

The session consisted of 209 trials broken into 7 blocks, where each block consisted of 30 trials (Fig. 2A). A two-way repeated measures ANOVA approach revealed a significant trial [F (2.972, 62.42) = 12.88, *P* < 0.0001] and treatment effect [F (2.691, 56.51) = 2.942, *P* < 0.05] on impulsivity. Dunnett tests revealed a significant decrease in number of baseline premature responses in fourth (*P*< 0.05), sixth (*P*< 0.01) or seventh (*P*< 0.01) trial block compared with first trial block, indicative of improvement in waiting impulsivity over time. Dunnett tests also showed that the highest dose of GBR increased waiting impulsivity in the second (*P*< 0.05), fourth (*P*< 0.05) and sixth (*P*< 0.01) trial block compared with control of the same block. The intermediate dose of GBR increased waiting impulsivity in the fifth trial block (*P*< 0.05) when compared with control of the same block. The gradual decrease in baseline waiting impulsivity as the session progresses could be due to change in motivation (Responsivity index), therefore, waiting impulsivity across the session was compared with responsivity across the session. Effects of GBR on responsivity index (motivation) over the course of the session is shown in Fig. 2B. A two-way repeated measures ANOVA approach revealed a significant trial effect [F (6, 592) = 3.228, *P* < 0.01] and no significant treatment effect [F (3, 592) = 1.973, *P* = 0.116] on responsivity index. Decrease in baseline responsivity index is indicative of waning motivation over the course of the session. Dunnett tests revealed a significant decrease in responsivity of 0.5 mg/kg treated rats in sixth trial block compared with 0.5 mg/kg GBR treated rats in the first trial block (*P*< 0.05), indicative of waning motivation. By comparing pattern between baseline premature responses and baseline responsivity index, the decrease in baseline premature response at the end of the session (trial block 7) can be determined to result not from waning motivation, but from learning mechanism (improvement in inhibitory response control over the course of the session).

**Figure 2.**
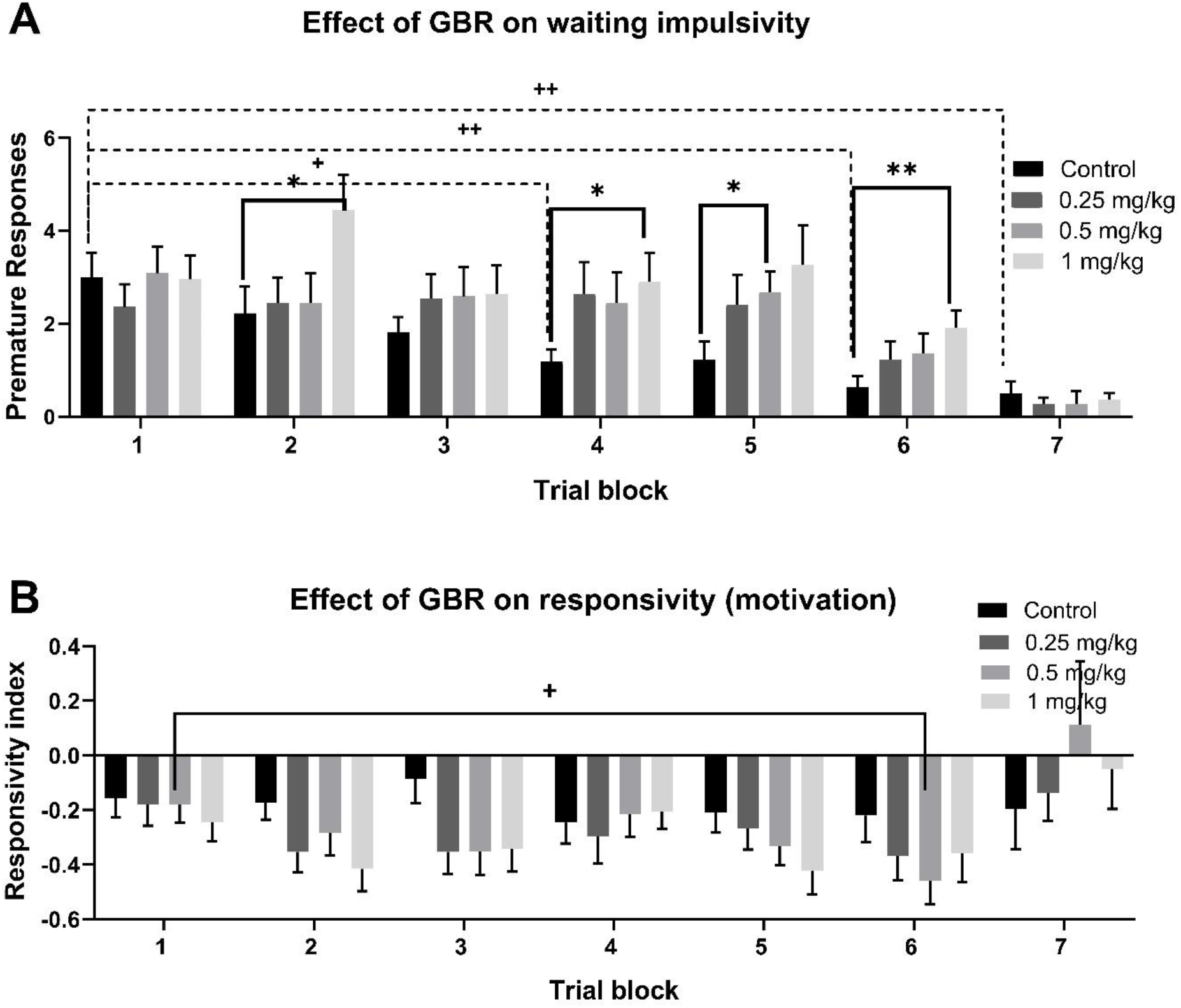
Effects of GBR 12909 (vehicle, 0.25, 0.5 or 1 mg/kg) on waiting impulsivity or responsivity index across the session (n=22). (A) Effects of GBR on waiting impulsivity across the session. (B) Effects of GBR on responsivity index across the session. Measures are shown as observed mean ±SEM. +*p* < 0.05 and ++ *p* < 0.01 compared to vehicle-treated rats (control) or GBR treated rats of 1st trial block. * *p* < 0.05 and ** *p* < 0.01, compared to control of the same block (Dunnett).

### Effect of GBR12909 on selective attention (accuracy) and omission

Main effect of GBR 12909 on attentional accuracy and omission averaged across different ITI is shown in Fig. 3A. Repeated measures one-way ANOVA revealed no significant treatment effect on accuracy [F (2.109, 44.29) = 2.713, P=0.07] and omission [F (2.741, 57.56) = 1.03, P=0.38]. Variability in baseline attentional performance was observed in individual animals and was used to divide rats into three subgroups. Rats were split into high-attentive (HA), mid attentive (MA) and low-attentive (LA) subgroups based on accuracy and number of misses made after saline administration (Fig. 3B). Previous studies report no significant change in the performance of mid attentive rats following drug administration (Hayward et al., 2016). Hayward et al. (2016) used an upper lower quartile split which included excluding the mid attentive rats. The exclusion of mid attentive rats resulted in a larger separation between HA and LA subgroups. Therefore, in this study the mid attentive rats were excluded from the analysis. Only the HA (accuracy > 95% and omission < 50%) and LA rats (accuracy < 80% and omission > 50%) were included in the analysis. Effects of GBR 12909 on accuracy and omission of HA and LA rats are shown in Fig 3B. A repeated measure two-way ANOVA revealed a Group x Treatment interaction for accuracy [F (3,33) = 9.782, *p*< 0.0001]. Accuracy was significantly higher in vehicle HA rats compared with vehicle LA rats (*p*< 0.0001). Dunnett’s test revealed that GBR significantly increased accuracy of LA rats at 0.25 and 0.5 mg/kg (all *p*<0.01) compared with vehicle control. GBR also reduced accuracy of HA rats at 0.25 mg/kg compared with vehicle control (*p*<0.05). A repeated measure two-way ANOVA revealed no Group x Treatment interaction for omission [F (3, 33) = 0.8443, P=0.47]. Re-analysis of the data using Oldham method revealed that the magnitude of improvement following administration of 0.25mg/kg and 0.5mg/kg GBR12909 was dependent on baseline accuracy (correlation analysis is shown in Fig. 3C). Oldham method’ revealed a strong negative association between baseline accuracy and the change following 0.25 mg/kg GBR and a moderate negative association between baseline accuracy and the change following 0.5 mg/kg GBR. Improved accuracy was seen in 7 out of 22 rats following 0.25 and 0.5 mg/kg GBR; rats with baseline accuracy < 85 % benefited the most from this treatment.

**Figure 3.**
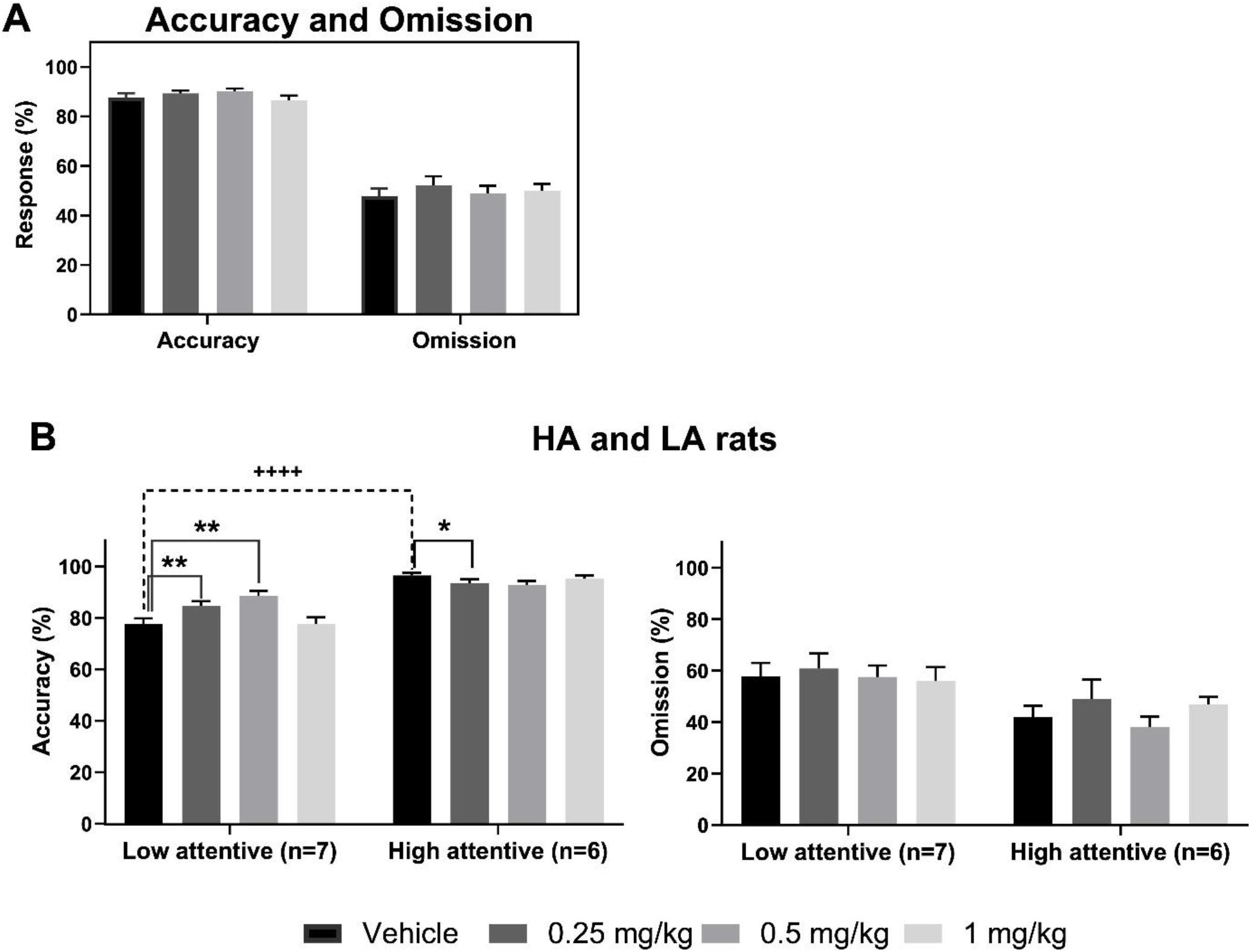
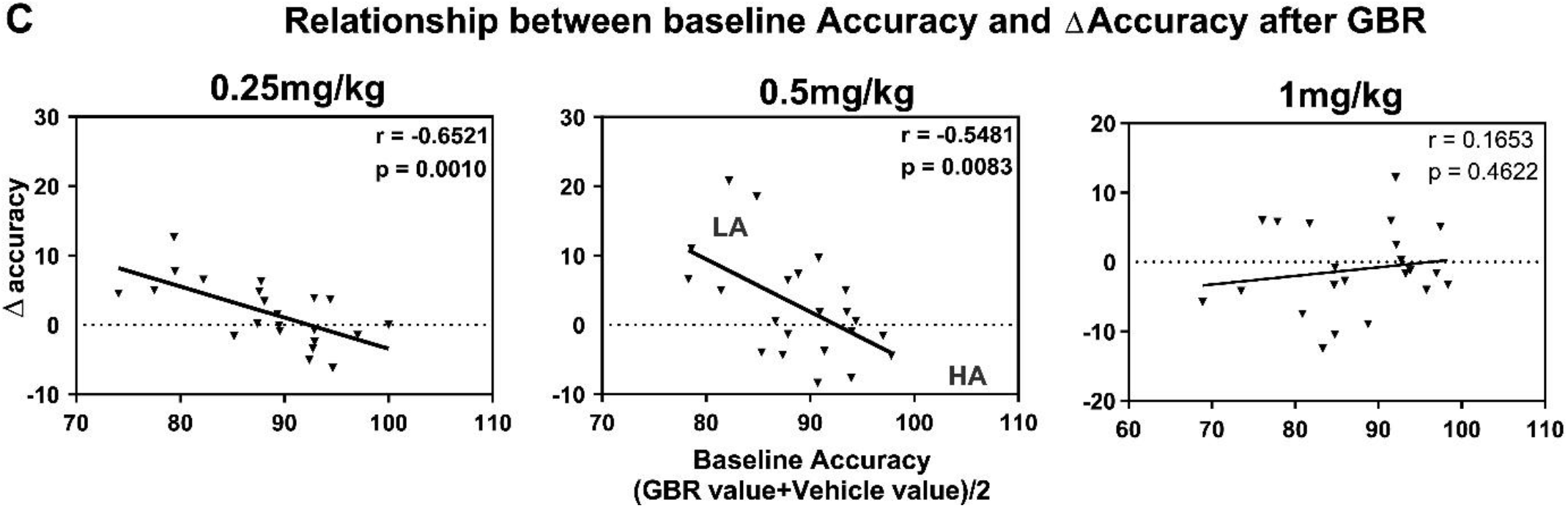
Effects of GBR12909 (vehicle, 0.25, 0.5 or 1 mg/kg) on performance accuracy and omission of rats in the 5C-CPT (n=22). (A) Group performance. (B) Subgroup performance for LA and HA rats. Mid attentive rats were excluded from the analysis. (C) Correlational analysis using Oldham method. Measures are shown as observed mean ±SEM. \**p* < 0.05, ** *p* < 0.01 compared to vehicle-treated rats (Control) of the same group. ++++*p* < 0.0001 compared to control of LA rats (Dunnett).

### Effects of GBR on compulsive behaviour

The main effect of GBR on compulsive behaviour is shown in Fig. 4 (A-C). GBR showed a tendency to decrease perseverative responding during target trials, although this did not reach significance [F (1.111, 23.33) = 1.088, *P*=0.3153]. In addition, GBR produced no significant change in perseverative responding when the number of perseverative responses during target and non-target trials was combined [F (1.239, 26.01) = 1.031, *P*=0.3364]. The highest dose of GBR showed a tendency toward reducing compulsivity in a manner consistent with baseline performance, however, it did not reach a significant level (Fig 4). Grubbs’ test did not detect an outlier. The relationship between baseline compulsivity and the drug-induced change was not affected by an outlier. Relationship between baseline Perseverative correct response or Perseverative false alarm and magnitude of change following administration of GBR (0.25, 0.5 or 1 mg/kg) is shown in Supplement Table 4.

**Figure 4.**
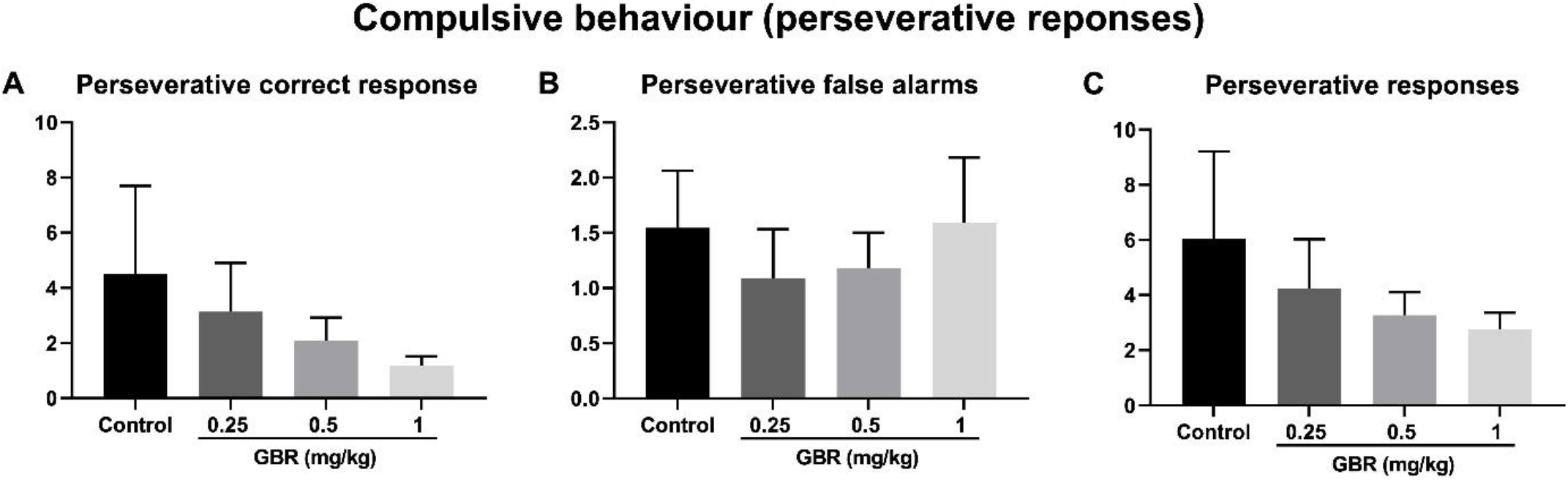
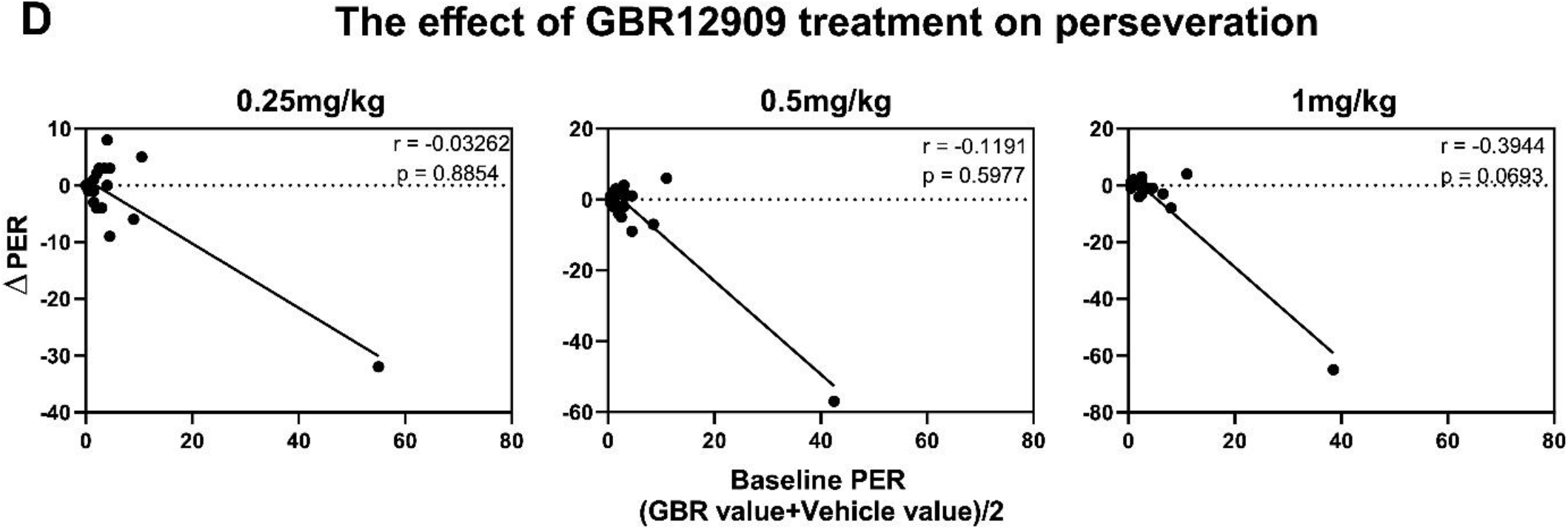
Effects of GBR (0.25, 0.5 or 1 mg/kg) on perseverative responding in the 5C-CPT. Effects of GBR on (A) perseverative correct responses, (B) perseverative false alarms, and (C) total number of perseverative responses. (D) Relationship between baseline compulsivity and magnitude of improvement following administration GBR. PER= perseverative responses.

### Measurements of Oldham’s angles

Angle measurements are shown in Fig 5. Amount of space between the regression lines were measured using a protractor. Three angles were measured: A, B and C as shown in Fig 5. Oldham’s angle measurements for the relationship between baseline accuracy and GBR induced change are shown in Fig 5A. A is the space between 1 mg/kg and 0.25 mg/kg regression lines. B is the space between 1 mg/kg and 0.5 mg/kg regression lines. C is the space between 0.5 mg/kg and 0.25 mg/kg lines (Fig 5A). If two angles are known, then the third angle can be calculated. For example, B=A+C. The measurement of the angle and the direction of movement (clockwise or counterclockwise) shows how effective is one dose at increasing accuracy of low attentive rats and reducing accuracy of high attentive rats compared with other doses. Oldham’s angle measurements for the relationship between baseline perseverative responses and GBR induced change are shown in Fig 5B-C. A is the space between the highest and the lowest dose (1 mg/kg and 0.25 mg/kg regression lines). B is the space between the highest and the intermediate dose (1 mg/kg and 0.5 mg/kg regression lines). C is the space between 0.5 mg/kg and 0.25 mg/kg lines (Fig 5B). If two angles are known, then the third angle can be calculated. A=B+C (Fig 5B-C). The clockwise movement (⟳) of the regression line as the dose increases shows that the highest dose is more effective than the lowest or the intermediate dose at reducing compulsive behaviour.

**Figure 5:**
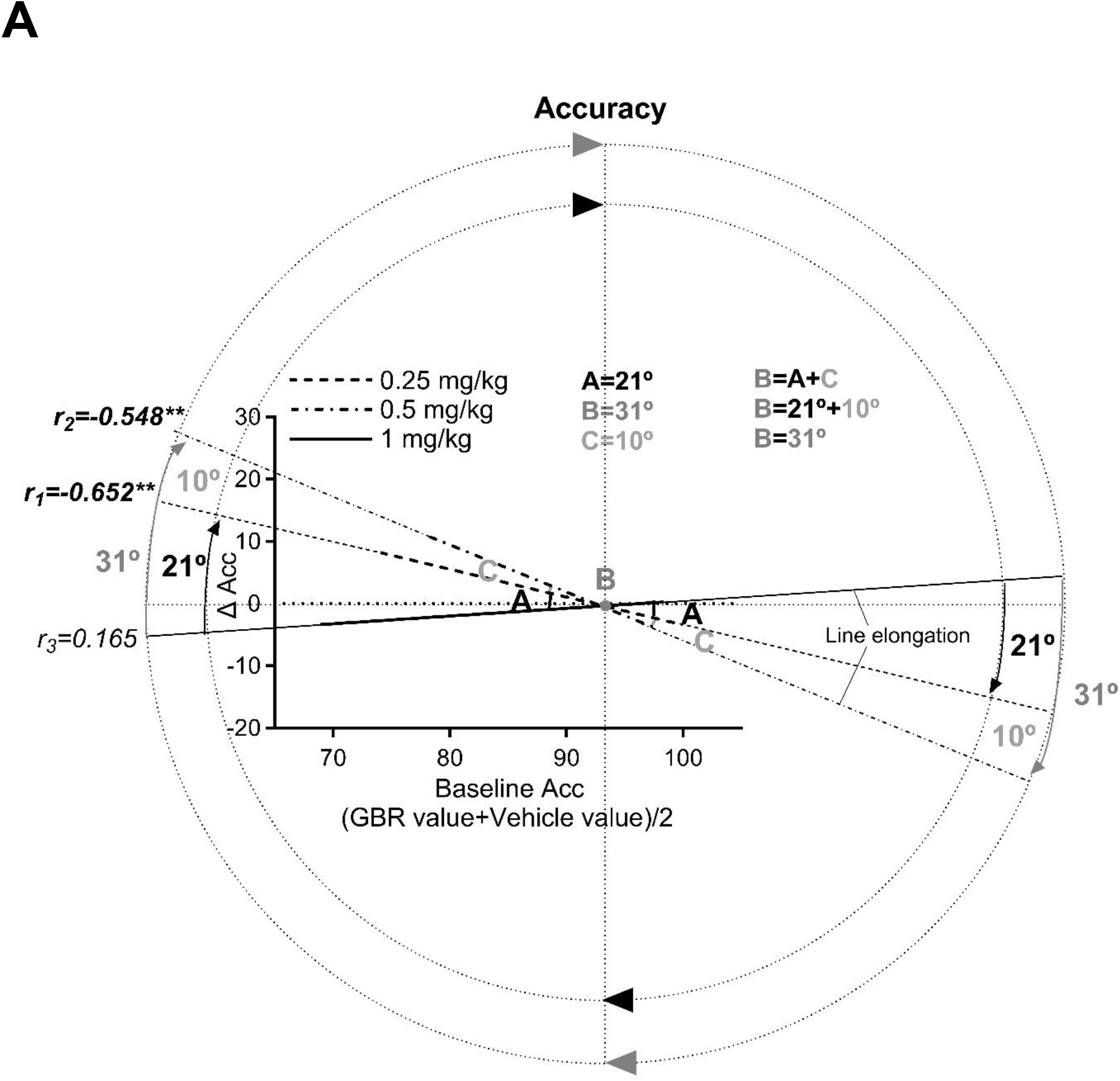
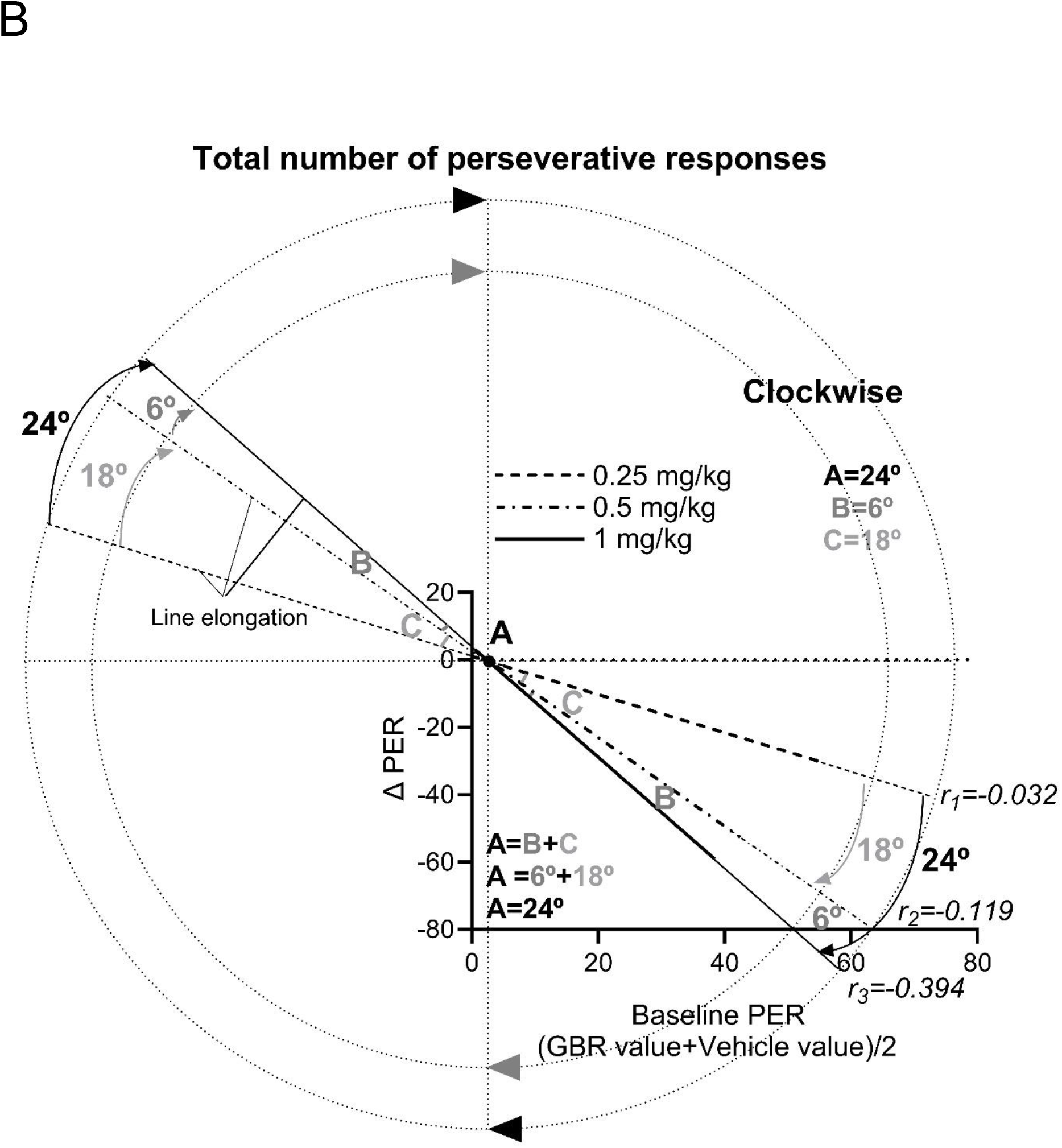
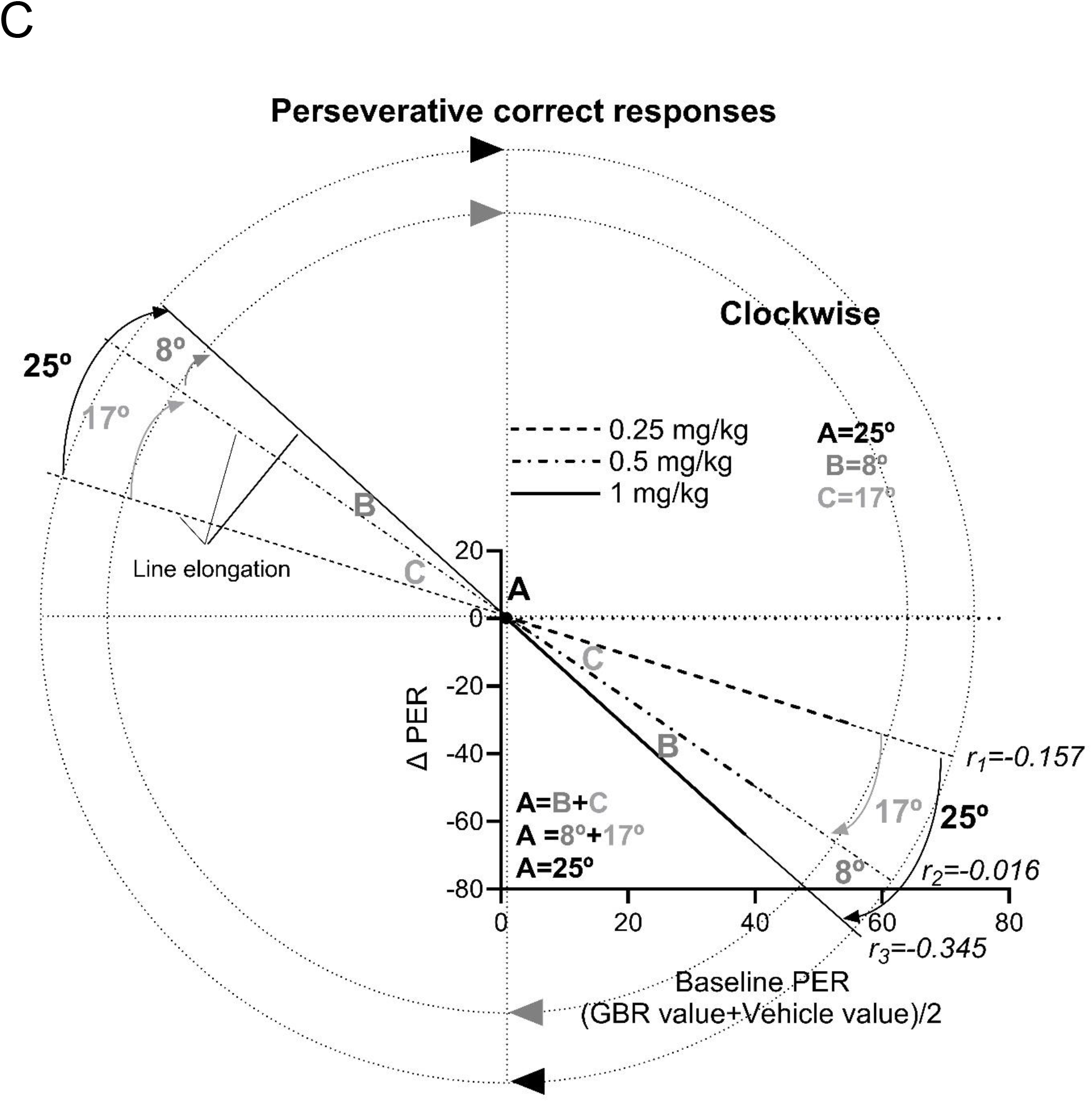
Measurement of Oldham’s angles. (A) Correlation between baseline accuracy and GBR induced change. (B) Correlation between baseline perseverative responses and GBR induced change. (C) Correlation between baseline perseverative correct responses and GBR induced change. Arrows show the clockwise movement of the regression line. The angles were measured using a protractor.

## Discussion

The current study highlights a preclinical approach to help address the pressing need for new ADHD medications that lack the treatment dependence issues seen with currently prescribed medications. Here, we examined the effects of the DAT antagonist GBR 12909 on 5C-CPT performance measures using the Oldham method in female Lister Hooded rats. Examination of the effect of this compound showed that the selective inhibition of DAT changes 5C-CPT performance measures in a rate-dependent manner. All doses of GBR changed waiting impulsivity in a baseline-dependent manner and the two lowest doses of GBR changed accuracy in a baseline-dependent manner. The highest dose of GBR showed a tendency toward changing compulsive behaviour in a rate-dependent manner. Taken together, the present results indicate that low doses of GBR improve accuracy of low-attentive rats and improve compulsivity in high-compulsive rats.

There is strong evidence to link DAT inhibition with inhibitory response control and visuospatial attention in rats (Volkow et al., 2007; Rubia et al., 2011; Dalley and Robbins, 2017). The selective DAT inhibitor GBR12909 has been consistently reported to increase premature responding in the 5-choice SRRT (5-CSRTT) (Baarendse et al., 2012; Fernando et al., 2012). For example, moderate doses of GBR12909 (2.5 and 5 mg/kg) impaired waiting impulsivity in high-impulsive rats; however, this impairment did not reach statistical significance (Fernando et al., 2012). In another 5-CSRT study, a higher dose of GBR12909 (10 mg/kg) caused a significant increase in the number of premature responses under standard and challenging test conditions (long ITI) (Baarendse et al., 2012). In the current study, systemic treatment with low doses of GBR12909 (0.25-1 mg/kg) impaired waiting impulsivity in a baseline dependent manner, further supporting a role for the dopamine transporter in regulating inhibitory response control as measured by 5C-CPT. The effect of GBR12909 on impulsivity is likely due to an increase in dopamine levels in subcortical regions, most likely the striatum (Dalley and Robbins, 2017). Correlation analysis using Oldham’s method revealed a direct relationship between baseline impulsivity and the change following GBR12909. A greater increase in impulsivity was observed in rats with high baseline impulsivity. The current study is the first to establish the presence of a rate-dependent relationship between baseline impulsivity and the change following GBR 12909. A within-session learning effect has previously been reported in studies using an extended ITI challenge with a constant SD (Young et al., 2011). By comparing patterns between waiting impulsivity and motivation (responsivity index), the decrease in baseline waiting impulsivity at the last trial block can be determined to result not from waning motivation, but from learning mechanism (improvement in inhibitory response control over the course of the session). Evidence for GBR-induced impairment in waiting impulsivity was observed when performance was examined across the session. GBR 12909 caused a dose dependent increase in waiting impulsivity, persisting until the end of the session. A statistically significant increase in waiting impulsivity following administration of the highest dose of GBR 12909 was observed after the first trial block. The impairment of waiting impulsivity was most substantial during the 6th trial block. This finding is consistent with previous studies reporting that systemic administration of GBR12909 (0.75 and 0.5 mg/kg) induces a significant DAT inhibition in rats within a matter of seconds, with dopamine uptake inhibition peaking 10-15 minutes post dosing (Espana et al., 2008).

Alongside the effect of GBR on waiting impulsivity, GBR (at 0.25 and 0.5 mg/kg) improved accuracy in low-attentive (LA) rats. Oldham’s method revealed an orderly relationship between baseline accuracy and response to GBR12909. In other words, the two lowest doses of GBR improved accuracy in rats with low baseline accuracy, but impaired accuracy in rats with high baseline accuracy, further supporting the inverted U function of dopaminergic activation in impulsivity (Tomlinson et al., 2015; Bickel et al., 2016). The pro-cognitive effect of GBR 12909 on attentional accuracy in the present study is consistent with the pro-cognitive effect of stimulants (which target DAT) in the signal detection task (Turner et al., 2017). The signal detection task is different from 5 choice tasks in that the rodent is required to detect a target visual stimulus presented repeatedly in the same location (central panel) rather than detecting a stimulus that may appear in any one of five locations (Turner et al., 2016; 2017). Recently the signal detection task has been used to examine the relationship between baseline accuracy and proportion of change in accuracy following amphetamine administration (Turner et al., 2016). Administration of a low dose of amphetamine improved attention by altering accuracy in low-attentive rats and this improvement correlated negatively with baseline accuracy performance (Turner et al., 2016; 2017). Correlational analysis using the re-test reliability method revealed a moderate negative association between baseline accuracy and the change following administration of amphetamine (Turner et al., 2017). These data demonstrated an increase in low baseline accuracy and a decrease in high baseline accuracy (Full range effect). Therefore, the effect of pharmacological manipulation on accuracy is dependent on heterogeneity of the performance prior to manipulation (baseline performance) (Bickel et al., 2016). This study further confirmed the importance of striatal dopamine in attentional processes by suggesting that the attentional impairment observed in low-performing rats could be due to dysregulation of the striatal dopaminergic system (Turner et al., 2017). However, the signal detection task is limited in that it does not include non-target trials, therefore, it cannot be used to measure response inhibition and sustained attention in a manner consistent with human CPT (Turner et al., 2016; 2017).The pro-cognitive effect of GBR 12909 on attentional accuracy in the present study is consistent with pro-cognitive effect of stimulants (which target DAT) in clinical studies (Koelega., 1993; Riccio et al., 2001). This is the first study to report the pro-cognitive effect of low doses of GBR, which was not revealed in the study by Fernando et al. (2012). The observation of such an effect here is likely because previous studies did not examine the relationship between baseline performance measures and the change following GBR12909 administration. Thus, correlational analysis used in this study were vital in revealing a relationship that would otherwise have gone unnoticed.

Dopaminergic neurotransmission in the NAcb has also been associated with compulsive drug seeking behaviour (Sanchez-Roige et al., 2012). In the present study, the highest dose of GBR 12909 showed a tendency toward improving perseverative correct response and total number of perseverative responses in a baseline dependent manner. This is the first study to report the presence of a negative relationship between baseline compulsivity and magnitude of change following administration of GBR 12909, suggesting that baseline levels of compulsive behaviour may influence the GBR 12909 effect. Thus, rats with high baseline compulsivity benefited the most from the treatment. This finding is consistent with previous studies in mice performing the Iowa gambling task (IGT), which report a decrease in perseverative responses following administration of GBR 12909 (Van Enkhuizen, et al., 2013). In addition, the abuse potential of amphetamine has been linked with its action on the mesolimbic dopaminergic pathway and it has been reported that GBR 12909 may be effective for the treatment of stimulant addiction (Ahn et al., 2013). The present study is also consistent with previous clinical studies that report that GBR 12909 (Vanoxerine) can be used for treatment of cocaine-dependent patients (Kadric et al., 2019). Subjects using cocaine are at increased risk of obsessive-compulsive disorder (OCD) (Crum et al., 1993) and there is an elevated risk of developing substance abuse in subjects with existing OCD (Rowe et al., 2022). As the main symptoms of OCD are perseveration and indecision (Miodownik et al., 2015), OCD is more prevalent among females in adolescence and adulthood (Mathes et al., 2019) and 40% of patients with OCD do not respond to current medications (Miodownik et al., 2015) the clinical investigation of novel analogues of GBR may well facilitate the discovery of new therapeutic agents for OCD. Future work can expand the current study, for example, 5C-CPT has been reverse translated to human, enabling compulsive behaviour to be studied in both OCD patients and rodents (Young et al., 2013). Together, these findings suggest that 5C-CPT can be used to measure impulsive and compulsive behaviour and these behaviours are independent.

In conclusion, the two lowest doses of GBR improved accuracy in female rats with low baseline accuracy, but impaired accuracy in female rats with high baseline accuracy, further supporting the inverted U function of dopaminergic activation in impulsivity. The highest dose of GBR 12909 showed a tendency toward improving compulsive behaviour in a baseline dependent manner. Rats with high baseline compulsivity benefited the most from the treatment. Unlike effects of GBR 12909 on selective attention and compulsive behaviour, GBR impaired impulsive behaviour in a baseline dependent manner. This study also highlights the value of Oldham’s method in revealing baseline-dependent effects of drugs on inhibitory response control and cognitive function in 5C-CPT. This study suggests that pre-drug baseline performance can influence the change from baseline following administration of GBR 12909 in rats. Novel analogues of GBR-type dopamine uptake blockers (GBR 12909 and GBR 12935) may have clinical utility for the treatment of obsessive-compulsive disorder, obsessive drug seeking behaviour, substance abuse in subject with existing OCD and inattentive subtype of ADHD (more prevalent among girls and adults).

